# Sequential effect and temporal orienting in pre-stimulus oculomotor inhibition

**DOI:** 10.1101/2023.04.04.535537

**Authors:** Noam Tal-Perry, Shlomit Yuval-Greenberg

## Abstract

When faced with unfamiliar circumstances, we often turn to our past experiences with similar situations to shape our expectations. This results in the well-established sequential effect, in which previous trials influence the expectations of the current trial. Studies have revealed that, in addition to the classical behavioral metrics, the inhibition of eye movement could be used as a biomarker to study temporal expectations. This pre-stimulus oculomotor inhibition is found a few hundred milliseconds prior to predictable events, with a stronger inhibition for predictable than unpredictable events. The phenomenon has been found to occur in various temporal structures, such as rhythms, cue-association and conditional probability, yet it is still unknown whether it reflects local sequential information of the previous trial. To explore this, we examined the relationship between the sequential effect and the pre-stimulus oculomotor inhibition. Our results (N=40) revealed that inhibition was weaker when the previous trial was longer than the current trial, in line with findings of behavioral metrics. These findings indicate that the pre-stimulus oculomotor inhibition covaries with expectation based on local sequential information, demonstrating the tight connection between this phenomenon and expectation and providing a novel measurement for studying sequential effects in temporal expectation.

## Introduction

Our day-to-day life is full of temporal uncertainty. We find ourselves asking, when will the next bus arrive? How long do I need to stand in the queue? When is my manuscript expected to be back from review? To guide our behavior through this uncertainty, e.g., to decide whether we should take a cab, switch to another queue, or contact our editor, we form temporal predictions regarding the likely onset time of expected events (Nobre & van Ede, 2018). The formation of temporal expectations can be based on different sources of information, i.e., temporal priors, such as associations with an informative cue (Amit et al., 2019; Coull et al., 2000; Miniussi et al., 1999) or prior statistical knowledge (Tal-Perry & Yuval-Greenberg, 2022). One such source of information is our recent experience, acquired through previous recent encounters with similar events. This form of temporal expectation is often called *sequential effects* – the expectation that the timing of the present event will resemble that of the previous one.

Sequential effects in the temporal domain are mostly studied using reaction time (RT) measurements (Capizzi et al., 2015; Possamai et al., 1973; Steinborn & Langner, 2012; Tal-Perry & Yuval-Greenberg, 2022). The typical trial design of these studies includes a warning signal followed, after a varying interval called *a foreperiod*, with a target. A common finding in this field is that when the foreperiod of one trial is shorter than its precedent, perceptual decisions regarding the present target will be slower. This was interpreted as reflecting the expectation that the duration of the present foreperiod will resemble that of the previous one. However, the opposite effect does not always occur, i.e., when the foreperiod of the present trial is longer than the precedent trial is, RTs remain unchanged relative to when the two trials have the same foreperiod (Niemi & Näätänen, 1981; Tal-Perry & Yuval-Greenberg, 2022).

While RT provides a useful behavioral measurement of temporal expectation, it measures temporal expectations, by definition, retrospectively, after the event has already occurred and expectations have already been formed. In a series of studies, we have shown that saccade rate can be used as a marker of temporal expectation, termed *the pre-target oculomotor inhibition effect*, which can be measured while expectations are formed rather than retrospectively. The pre-target oculomotor inhibition effect is a reduction in the rate of eye movements which occurs a few hundred milliseconds prior to the appearance of a predictable, relative to an unpredictable, target (Abeles et al., 2020; Amit et al., 2019; Badde et al., 2020; Dankner et al., 2017; Tal-Perry & Yuval-Greenberg, 2020, 2021).

The inhibition of eye movements during the foreperiod joins a host of other evidence pointing to the crucial role inhibition plays in many forms of temporal expectation, including sequential effects (Los, 2013). Transcranial magnetic stimulation (TMS) studies in human participants have shown that during motor response preparation, the motor-evoked potentials of both task-relavent and task-irrelavent muscles are inhibited prior to the anticipated target onset, indicating that corticospinal excitability is suppressed in order to prevent premature response (Duque & Ivry, 2009), which functionally may help to improve the signal-to-noise ratio at the moment a response is required (Greenhouse et al., 2015). Invasive measures in animals likewise indicate a pre-movement global inhibition of the musculatory system (e.g., Prut & Fetz, 1999), and it is evident that interfering with these inhibitory processes impairs timely response execution (Narayanan et al., 2006). As previously discussed (Tal-Perry & Yuval-Greenberg, 2021), this preparatory global inhibition of the motor system may spread to the oculomotor system, resulting in the inhibition of eye movements prior to expected stimuli.

The oculomotor inhibition effect was shown to be modulated by various types of temporal expectations, including those driven by rhythm (Dankner et al., 2017), by associations between a cue and a target, and by hazard rate – the change in the conditional probability for target occurrence that changes when times passes by and the target has not yet appeared (Amit et al., 2019; Tal-Perry & Yuval-Greenberg, 2020). However, to date, it remains unknown whether the pre-stimulus oculomotor inhibition marker is modulated solely by global temporal orienting or also by local sequential information, i.e., temporal predictions induced by the recent previous trials. Finding a link between sequential effects and the oculomotor inhibition effect would provide supporting evidence for a link between oculomotor inhibition and the preparatory inhibition observed in previous studies. Furthermore, it would indicate that this marker is associated with the formation of temporal expectations based on recent previous experiences. Such a finding would lend further support to the validity of this index as a measurement of temporal expectations and provide a novel metric for studying the effect of local sequential information.

In this study, we examined the links between oculomotor inhibition, the sequential effects, and temporal orienting in two experiments. In Experiment 1 (N=40), we examined saccade rate as a function of the difference between the foreperiod of one trial and that of its precedent trial. We hypothesized that local sequential information is reflected in eye movements dynamics – that when the foreperiod of a given trial is shorter than that of its precedent, there would be less pre-target oculomotor inhibition (i.e., more pre-target saccades) relative to when it was longer or equal. Our findings confirmed this hypothesis, with the sequential effect on saccade inhibition paralleling the pattern usually reported for RTs in previous studies. In Experiment 2 (*N*=40) we examined whether this first-order oculomotor sequential effect could provide an alternative interpretation to our previous findings of the pre-target oculomotor inhibition effect. This study joins previous studies by showing that the pre-stimulus oculomotor inhibition covaries with temporal expectation of various temporal priors and modalities, supporting its validity as an index for the formation of temporal expectations. Both experiments include new analyses of previously published datasets (Amit et al., 2019; Tal-Perry & Yuval-Greenberg, 2020, 2022). The previous studies either did not include an eye tracking analysis (Experiment 1) or did not include analysis of sequential effects (Experiment 2). Notably, here, as in our previous studies on the oculomotor inhibition effect (Abeles et al., 2020; Amit et al., 2019; Tal-Perry & Yuval-Greenberg, 2020, 2021), we use the general term “saccades” to include both large saccades and miniature saccades performed during fixation, which fit the definition of ‘microsaccades’ (Martinez-Conde et al., 2004, 2009). With this decision, we rely on the common view that saccades and microsaccades constitute an oculomotor continuum, both activated by similar neural mechanisms and sharing similar functions (Otero-Millan et al., 2008, 2013). The findings are similar when only small saccades (<1 visual degree) are included (see **Supplementary Material S1**).

## Methods

### Participants

A total of 80 participants were included in the study: 40 participants were included in Experiment 1 (25 females, 3 left-handed, Mean age 24.95±3.82 standard deviations [SD]) and 40 participants in Experiment 2 (24 females, one left-handed, Mean age 22.55±3.12 SD). All participants were healthy, reported normal or corrected-to-normal vision, and reported no history of neurological or psychiatric disorders. Participants received payment or course credit for their participation. The experimental protocols were approved by the ethics committees of Tel-Aviv University and the School of Psychological Sciences. Prior to participation, participants signed informed consent forms. The present study includes two experiments that are based on reanalyses of three published data sets as follows: Exp. 1 consisted of a novel analysis of eye tracking data of participants who were originally included in Tal-Perry & Yuval-Greenberg (2022), but no eye tracking analysis was performed in the original study; Exp. 2 consisted of a reanalysis of the eye tracking data of datasets published originally in Amit et al. (2019) and Tal-Perry & Yuval-Greenberg (2020).

### Stimuli

#### Experiment 1

The fixation object consisted of a dot (0.075° radius) within a ring (0.15° radius), embedded within a diamond shape (0.4×0.4°). The edges of the diamond changed color from black to white, cueing attention to the left (two left edges became white) or right (two right edges became white) side of the fixation object, or remaining neutral with respect to the target location (all four edges became white). The target was a black asterisk (0.4×0.4°) presented at 4° eccentricity to the right or left of the fixation object. A 1000 Hz pure tone was played for 60 ms as negative feedback following errors. Fixation objects and targets were presented on a mid-gray background.

#### Experiment 2

The fixation object in this experiment was a cross (black or blue, 0.4° × 0.4°), and the target was a Gabor grating patch (2° diameter, 30% contrast, spatial frequency of 5 cycles/degree) slightly tilted clockwise (CW) or counter-clockwise (CCW) from vertical, with tilt degree determined individually via 1-up 3-down staircase procedure. All stimuli were displayed at screen center on a mid-gray background.

### Procedure

The datasets that we have used originated from three previously published studies. Exp. 1 was based on the dataset reported in Tal-Perry & Yuval-Greenberg (2022). Exp. 2 was based on the combined datasets of Amit et al., (2019) and part of the dataset of Tal-Perry & Yuval-Greenberg (2020), both based on identical procedures.

#### General procedure

In both experiments, participants were seated in a dimly lit room, with a computer monitor placed 100 cm in front of them (24" LCD ASUS VG248QE, 1,920 × 1,080 pixels resolution, 120 Hz refresh rate, mid-gray luminance was measured to be 110 cd/m^2^). During the session, participants rested their heads on a chinrest. MATLAB R2015a (Mathworks, USA) was used to code and control the experiment, with stimuli displayed using Psychophysics Toolbox v3 (Brainard, 1997).

#### Experiment 1

Each trial started with a central black fixation object, presented until an online gaze-contingent procedure verified 1000 ms of stable fixation, defined by the placement of gaze within a radius of 1.5° of screen center. Following this, the edges of the fixation object changed color for 200 ms to represent a spatial cue (right or left in 75% of trials or neutral in the remaining 25% of trials). After a varying foreperiod (500 / 900 / 1300 / 1700 / 2100 ms) the target was briefly (33 ms) presented at 4° eccentricity to the left or right of the screen center, with the cue being valid with the target location in 75% of informative trials. Participants were instructed to respond as fast as possible upon detecting the target via a single button press. Between groups, participants were presented with the five foreperiods in either a uniform distribution (20% probability for each foreperiod) or an inverse-U-shaped distribution (a ratio of 1:2:3:2:1 between the five foreperiods, leading to trial percentages of approximately 11%, 22%, 33%, 22%, and 11%, respectively). Fixation was monitored throughout the foreperiod, using an online gaze-contingent procedure, and trials that included ≥ 1.5° gaze-shift for more than 10 ms during this period were aborted and repeated at a later stage of the session. An error feedback tone was played when participants responded before the target onset or did not respond within 1000 ms following the target onset. These trials were not included in the analysis. Participants of the uniform distribution group (N=20) performed 10 blocks of 160 trials each, divided into two sessions. Participants of the inverse-U-shaped distribution group (N=20) performed 18 blocks of 144 trials each, divided into three sessions done on separate days. A short break was given after each block. A practice block of 10 trials with random conditions was administered at the beginning of each session. **Figure 1A** summarizes the trial procedure used in this experiment.

**Figure 1.**
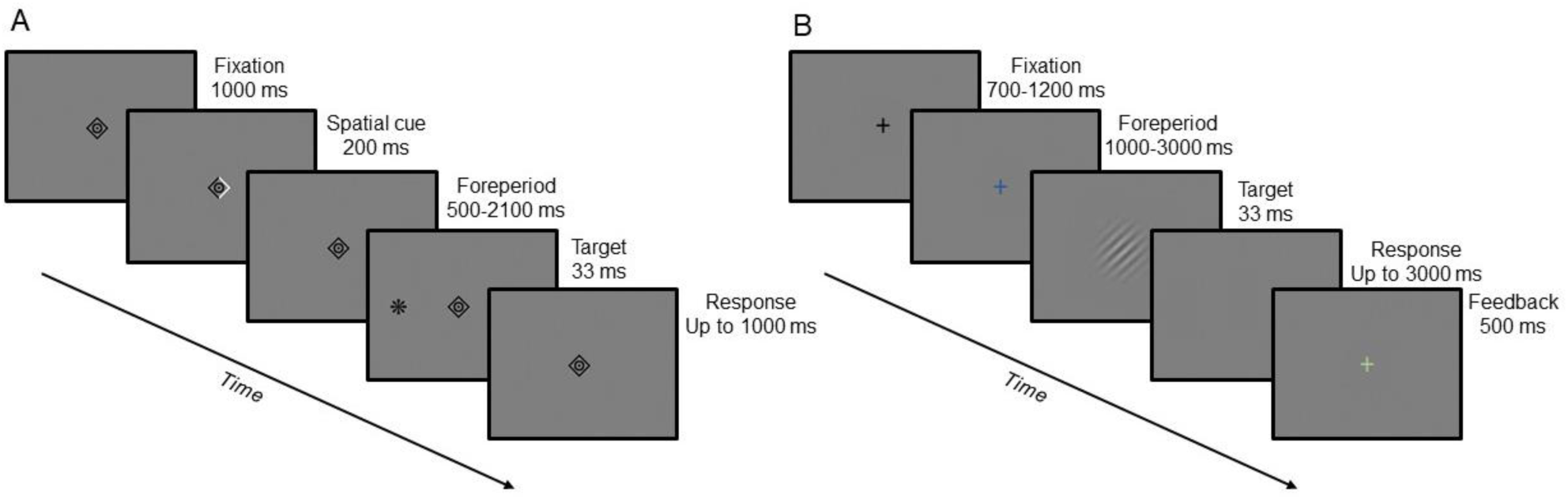
Trial progression. (A) Trial progression for Exp. 1, adapted from Tal-Perry & Yuval-Greenberg (2022); (B) Trial progression for Exp. 2, adapted from Tal-Perry & Yuval-Greenberg (2020), which was identical to the trial progression used in Amit et al. (2019).

#### Experiment 2

A central black fixation cross was presented between trials for a jittered inter-trial interval of 700-1200 ms. At trial onset, the fixation cross changed color from black to blue marking the onset of the foreperiod interval. After the foreperiod had elapsed, the target (tilted Gabor patch) was briefly (33 ms) presented and followed by a blank screen, and participants were requested to perform a 2AFC discrimination on the Gabor tilt by pressing one of two keyboard keys. Foreperiod was set to be between 1000-3000 ms in 500 ms increments. In fixed blocks, foreperiod was constant throughout the block. In random blocks, the foreperiod randomly varied from trial to trial, with foreperiods uniformly distributed. In part of the dataset (originally published in Amit et al., 2019), participants completed a total of five fixed blocks (one of each foreperiod) and five random blocks, with 100 trials per block. In the rest of the dataset (originally published in Tal-Perry & Yuval-Greenberg, 2020), participants completed two fixed blocks (one of 1000 ms and one of 2000 ms), and five random blocks, with 80 trials per block. Ten additional blocks performed by the participants in Tal-Perry & Yuval-Greenberg (five blocks with a distribution of 80% 1000 ms foreperiods and 20% 2000 ms foreperiods, and five blocks with the opposite ratio) were not included in the present study. The collapsed datasets of the two original studies were imbalanced in their number of trials per condition: there were 500 random trials and 500 fixed trials of five different foreperiods per participant in Amit et al. and there were 400 random trials and 160 fixed trials of two different foreperiods per participant in Tal-Perry & Yuval-Greenberg. This imbalance was accounted for by the generalized linear mixed model analysis which takes into account trial-wise variance (see Statistical analysis). **Figure 1B** summarizes the trial procedure used in this experiment.

### Eye-tracking

Eye movements were monitored using EyeLink 1000 Plus infrared video-oculographic desktop mounted system (SR Research Ltd., Oakville, ON, Canada), with a 1000 Hz sampling rate and the standard online and offline analog filters provided by Eyelink (Stampe, 1993). A nine-point calibration was performed at the beginning of the experiment and when necessary. Raw gaze data was low-pass filtered at 60 Hz and segmented between -500 ms relative to cue onset and 500 ms relative to target onset. Blinks were detected based on the built-in algorithm provided by EyeLink, plus an additional criterion requiring a binocular change in pupil size that exceeded 2.5 standard deviations from the segment’s mean pupil size for 3 or more consecutive samples (Hershman et al., 2018). Saccades were detected using a published algorithm (Engbert & Kliegl, 2003; Engbert & Mergenthaler, 2006), with saccade onset defined as the point in which the absolute standardized eye velocity exceeded the segment’s median eye velocity by six or more standard deviations, for a minimum of six consecutive samples. Only binocular saccades were included in the analysis. A 50 ms interval between saccade offset and the next saccade onset was imposed to prevent detection of overshoots. Intervals with blinks and 200 ms before the onset of blinks and after their offsets, were excluded from the saccades analysis. The analysis included saccades of all sizes, although most saccades (91.12%) were minuscule (<1°) due to the instruction to maintain fixation, and thus fit the definition of microsaccades (Martinez-Conde et al., 2009). Correlation between saccade amplitude and peak velocity (main sequence, Zuber et al., 1965) was high (*r* > 0.9) for all participants, verifying the validity of the saccade detection procedure.

For each trial, we determined whether a saccade was detected in the period of -300 to 0 ms relative to the target onset. This range was based on the time period known to include the oculomotor inhibition effect as reported in Tal-Perry & Yuval-Greenberg (2021). Trials that included a blink or missing data during this period were discarded from the analysis.

### Statistical analysis

#### Experiment 1

For each trial, we calculated the difference between its foreperiod and the foreperiod of the previous trial (FP-difference, *FP_n_* − *FP*_*n*−1_). For this purpose, the first trial of each session was discarded from the analysis. The resulting continuous factor ranged between - 1600 and 1600 ms in 400 ms increments and was Z-scaled to reduce computational complexity (standardized foreperiod difference, SFD). The probability of a saccade onset in the pre-target time interval was then analyzed using a generalized linear mixed model (GLMM), assuming a binomial family of responses (saccade present / absent) with a logit link, i.e., a logistic mixed model. We based our choice of the model on the assumption that due to pre-stimulus oculomotor inhibition, in the vast majority of trials, we are not expecting more than a single saccade to occur in the analyzed duration, with trials deviating from this assumption being rare and of little impact on the overall results. The following fixed factors were included in the model: (1) a scaled linear and quadratic relation between the current and previous foreperiod (SFD), to model the sequential effect; (2) the foreperiod distribution (uniform / inverse-U-shaped), a between-subject factor, using sum contrasts; (3) The interaction between the two factors, to model the effect of foreperiod distribution on the sequential effect. As oculomotor behavior was examined prior to target onset, the trial’s spatial cueing condition used in the original experiment (valid / invalid / neutral) was not included as a factor.

#### Experiment 2

Trials of the random block were screened according to the foreperiod of the previous trial, such that only trials in which the previous foreperiod was equal to the current trial’s foreperiod were included in the analysis. The probability of a saccade onset was then analyzed using a logistic mixed model, with the following factors: (1) the current trial’s foreperiod; (2) condition (fixed / random); and (3) the interaction between the two factors. Difference contrast was used for Foreperiod, and sum contrast was used for Condition.

#### General statistical analysis

For all models, statistical significance for main effects and interactions was determined via a likelihood-ratio (LR) test against a reduced nested model excluding the fixed term (i.e., type-II sum of squares, SS), and statistical significance for parameter coefficients was determined according to Wald z-test (Fox, 2016). To provide support for null results (*p* > 0.05), we additionally calculated the Bayes Factor (BF) between the full and reduced model, using BIC approximation (Wagenmakers, 2007). BF is reported with the null result in the denominator (*BF*_01_), representing how much the data is supported by the null model relative to the full model. The model’s random effect structure was selected according to the model that was found to be most parsimonious with the data, i.e., the fullest model that the data permits while still converging with no singular estimates (Bates, Kliegl, et al., 2015), in order to balance between type-I error and statistical power (Matuschek et al., 2017). This was achieved by starting with a random intercept by subject-only model and continuing to a model with random slopes for fixed terms by subject and their correlation parameters, and from there to a random-interaction-slopes by-subject model, testing for model convergences in each step. Models that failed to converge were trimmed by the random slope with the least explained variance and were retested. Analyses were performed in R v4.0.3 using R-studio v1.3.959 (R Core Team, 2018). Modelling was performed using the lme4 (Bates, Mächler, et al., 2015) package, BF was calculated using the BayesFactor package (Morey & Rouder, 2018), and model diagnostics were performed using the performance package (Lüdecke et al., 2020). An R-markdown file describing all the model fitting steps and diagnostic checks on the final model is available at the project’s OSF repository (see Data Availability Statement)

## Results

### Experiment 1: the oculomotor sequential effect

In the first experiment, we tested whether the pre-stimulus oculomotor inhibition is affected by contextual information about the previous trial in a speeded detection task. For this goal, we analyzed the eye movement data from Tal-Perry & Yuval-Greenberg (2022). We first start by examining the behavioral sequential effect in the same dataset.

#### Reaction times

Participants’ mean reaction times (RTs) were reported in a previous publication (Tal-Perry & Yuval-Greenberg, 2022) and are summarized here. After discarding trials with no previous foreperiod, trials with no response, and trials in which a response occurred within less than 150 ms relative to target onset, the RTs of the remaining trials were modeled as a factor of the Standardized Foreperiod Difference between the current and the preceding trial (SFD, continuous; linear and quadratic terms), Foreperiod Distribution (uniform / inverse-U-shaped), Condition (valid / invalid / neutral), and their interaction terms. Results for the valid condition are displayed in **Figure 2A**. Findings revealed a strong sequential effect, such that RT was slower in trials that were preceded by a longer trial, while the opposite was not found. This effect was modulated by the Foreperiod Distribution, but not by Condition. These results confirm that a behavioral sequential effect was present in the analyzed dataset.

**Figure 2.**
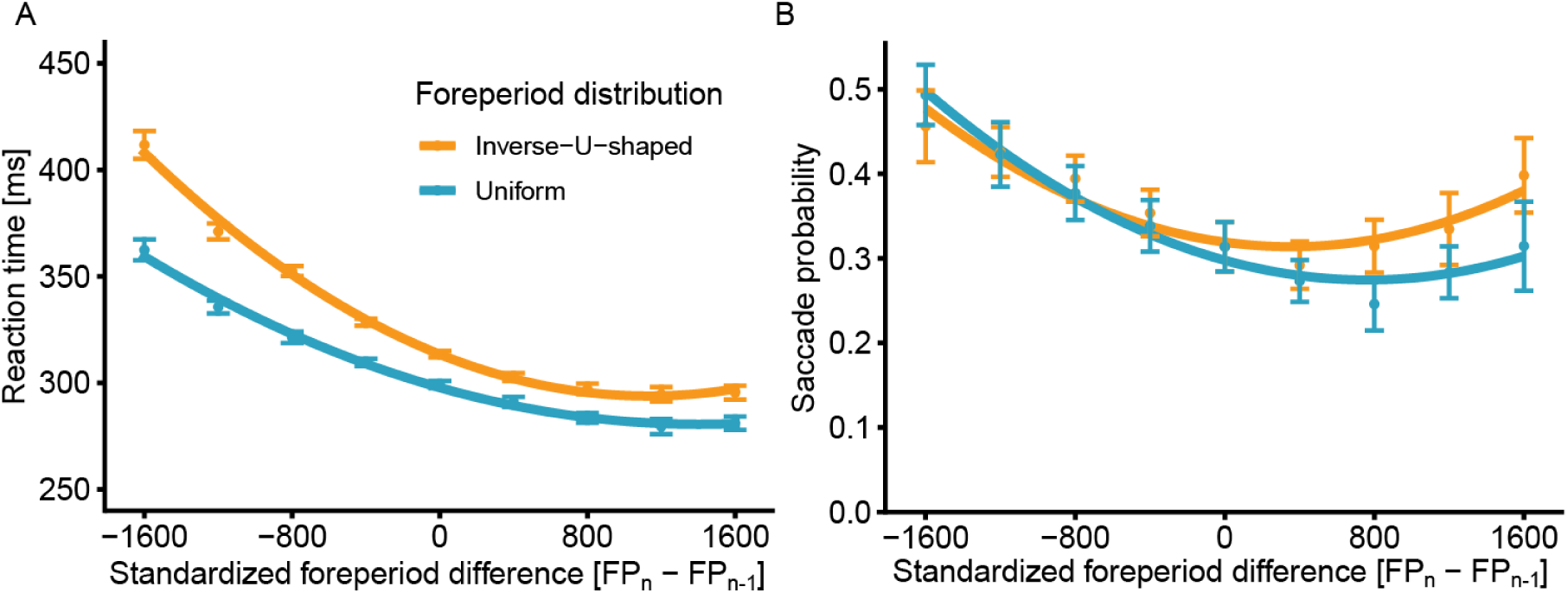
Experiment 1 results. Reaction time in valid cue trials (A) and the probability of performing a saccade during the -300 to 0 ms period relative to target onset (B), as a function of the difference between the current and previous foreperiod, and the Foreperiod Distribution. Negative values indicate that previous foreperiod was longer than current foreperiod, and vice-versa for positive values. Error bars depict ±1 standard error from the mean, correcting for within-subject variability (Cousineau & O’Brien, 2014). Lines depict 2^nd^ polynomial fit to the observed data. *N* = 20 in each distribution.

#### Eye tracking

The probability of a saccade occurring prior to target onset was examined using a logistic mixed model, with SFD (continuous; linear and quadratic terms), Foreperiod Distribution (uniform / inverse-U-shaped), and the interaction between them set as fixed factors, allowing for a random intercept per subject and a random slope to SFD per subject.

The saccade probability for each distribution as a factor of the standardized difference between the current and previous foreperiod is displayed in **Figure 2B**. Consistently with the RT findings, results showed that SFD had a strong influence on saccade likelihood (*χ*^2^(2) = 30.225, *p* < .001), with a strong negative linear slope (log estimate -0.238, *z* = -6.061, *p* < .001), indicating that saccade probability is reduced as the difference between the current and previous trial gradually turns from being negative (previous is longer) to positive (previous is shorter). This indicates that oculomotor inhibition is enhanced when the previous trial becomes shorter relative to the current trial. This negative slope was accompanied by a weaker positive quadratic slope (log estimate 0.081, *z* = 4.483, *p* < .001), reflecting that the degree of saccade inhibition is gradually decreased until it changes direction. The combination of the negative linear and the positive quadratic trends resulted in the asymmetry between negative and positive SFD observed.

To examine this gradual change, we modeled SFD as a categorical factor and contrasted each level with an SFD of zero (i.e., previous foreperiod equals current; see **Table 1**). As can be observed, the negative SFDs (longer foreperiod at previous trial) resulted in higher saccade probability compared to the zero SFD (no difference in foreperiod between this and the previous trial), with a gradual decrease in saccade probability difference as the SFD decreases. The positive SFDs (shorter foreperiod at previous trial) resulted in a significantly smaller saccade probability for the 400 and 800 ms difference relative to the zero SFD, with the effect gradually decreasing as the SFD increases, such that no significant difference was observed for the larger positive SFD values. These findings are consistent with the hypothesis that oculomotor inhibition reflects anticipation – in negative SFD trials, participants are anticipating the target to occur at a later stage of the trial, thus there is a weaker inhibition of saccades prior to the actual target onset as compared to trials where the target appeared at the expected time. Assuming the participants have learned the distribution of foreperiods, reduced saccade rate observed in the first two positive SFD levels may reflect the effect of conditional probability (hazard rate) – as target was expected to appear but has yet to appear, anticipation continues to be built up, and with it, the inhibition of saccades continues to increase; or it may reflect the aggregated effect of earlier foreperiods, i.e., higher-order sequential effects. Lastly, the lack of significant difference in the last two positive SFD levels may reflect the added time uncertainty in expectation – at these trials, target appeared long after the expected time, such that anticipating when would it occur became increasingly more difficult.

**Table 1.**
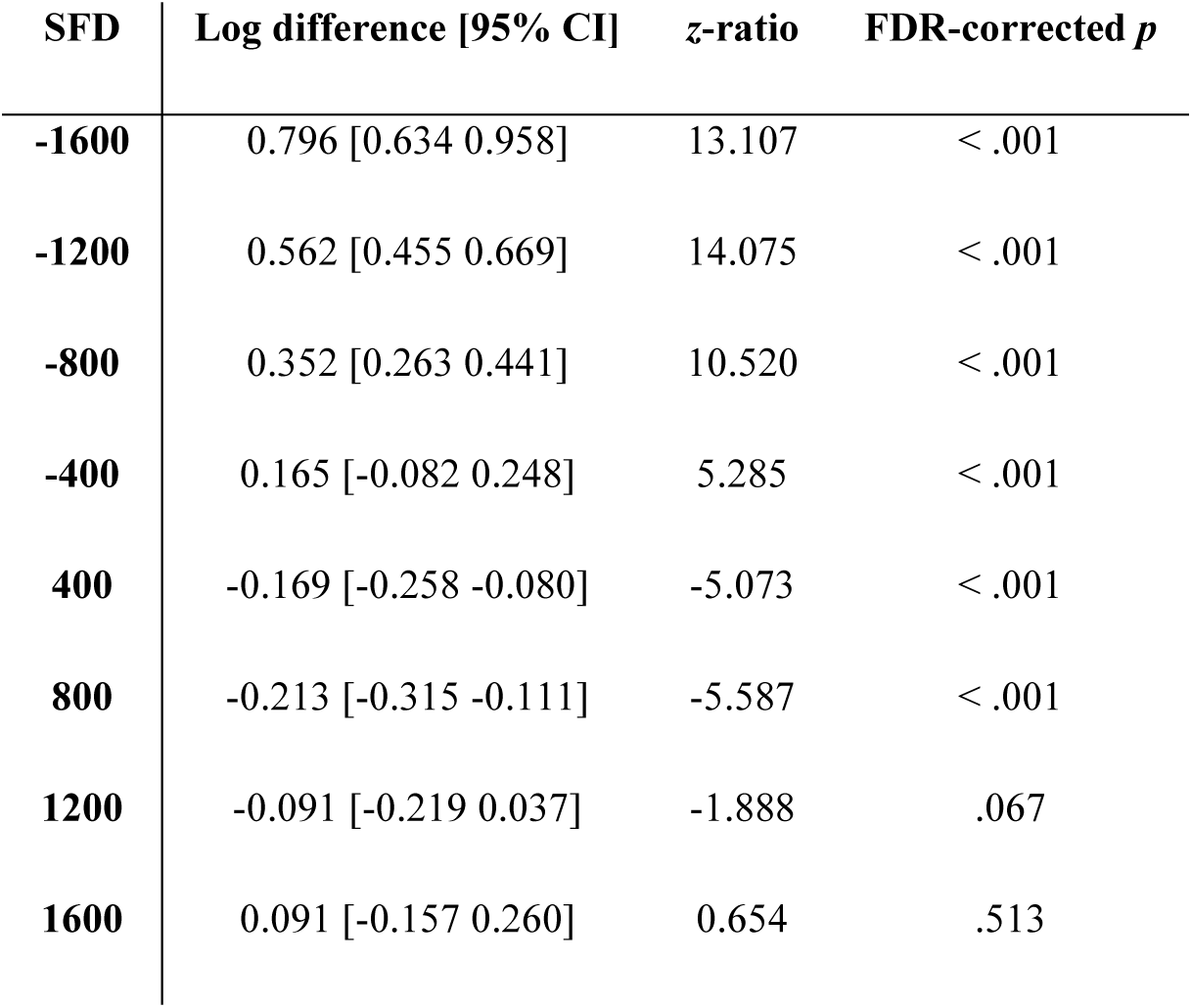
SFD from zero level. The model estimates represent the difference in log-odds saccade probability of [SFD level minus zero level] along with a 95% confidence interval (CI). Reported statistics based on log odds-ratio with the zero SFD level in the denominator. Reported *p* values were false discovery-rate (FDR) corrected. SFD = standardized foreperiod difference

Importantly, the SFD significantly interacted with the Foreperiod Distribution (*χ*^2^(2) = 17.635, *p* < .001), stemming from both a difference in the linear (log estimate 0.038, *z* = 3.529, *p* < .001) and quadratic (log estimate 0.022, *z* = 2.554, *p* = .011) components between the foreperiod distributions. To explore this interaction, we contrasted each SFD between distributions (see **Table 2**). As can be observed, the two foreperiod distributions significantly differed only for the positive SFD values starting with 800 ms. Interestingly, this pattern of interaction differed from the one observed in the RT data (**Figure 2A**), in which the greatest differences between the distributions were observed for the most negative SFDs, with differences decreasing as SFD got closer to zero and went into positive values. These diverging patterns may stem from underlying mechanistic differences between the motor and oculomotor systems. Lastly, we observed a significant effect for Foreperiod Distribution (*χ*^2^(1) = 23.588, *p* < .001), stemming from the higher probability of saccade occurrence for the inverse-U-shaped distribution. These results suggest that the pre-stimulus oculomotor inhibition reflects the combination of short-term expectations from the previous trials and long-term expectations from the distribution of intervals and higher-order sequential effects.

**Table 2.**
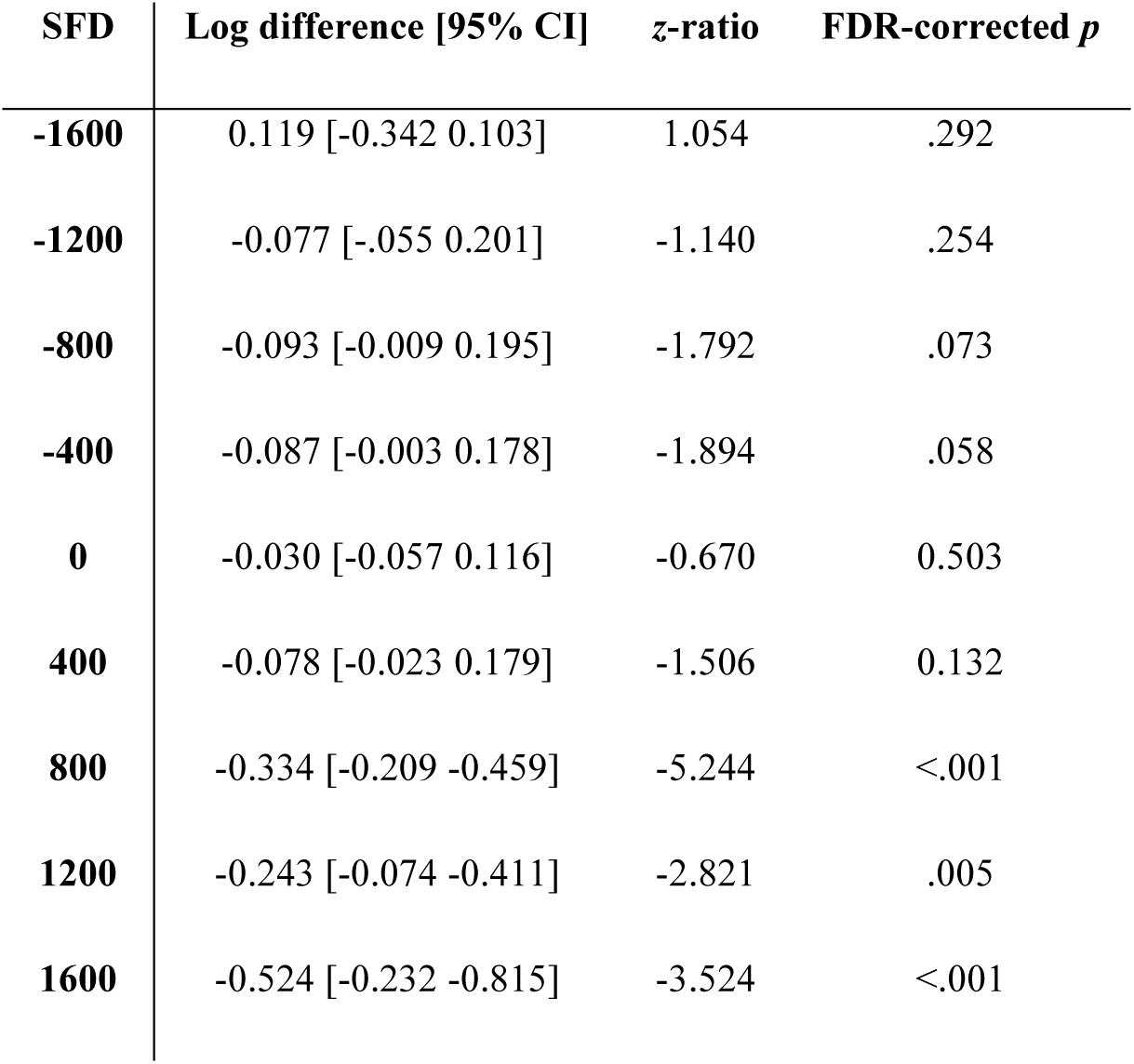
SFD contrasts by Foreperiod Distribution. The model estimates represent the difference in log-odds saccade probability [uniform minus inverse-U-shaped] conditions along with a 95% confidence interval (CI). Reported statistics based on log odds-ratio with uniform distribution in the nominator. Reported *p* values were false discovery-rate (FDR) corrected. SFD = standardized foreperiod difference.

| SFD | Log difference [95% CI] | z-ratio | FDR-corrected *p* |
| --- | --- | --- | --- |
| -1600 | 0.119 [-0.342 0.103] | 1.054 | .292 |
| -1200 | -0.077 [-.055 0.201] | -1.140 | .254 |
| -800 | -0.093 [-0.009 0.195] | -1.792 | .073 |
| -400 | -0.087 [-0.003 0.178] | -1.894 | .058 |
| 0 | -0.030 [-0.057 0.116] | -0.670 | 0.503 |
| 400 | -0.078 [-0.023 0.179] | -1.506 | 0.132 |
| 800 | -0.334 [-0.209 -0.459] | -5.244 | <.001 |
| 1200 | -0.243 [-0.074 -0.411] | -2.821 | .005 |
| 1600 | -0.524 [-0.232 -0.815] | -3.524 | <.001 |

### Experiment 2: Does the fixed effect stem from the sequential effect?

Previous studies have demonstrated that the pre-stimulus oculomotor inhibition is stronger when the foreperiod is fixed throughout the block compared to when it varies randomly (Abeles et al., 2020; Amit et al., 2019; Badde et al., 2020; Dankner et al., 2017; Tal-Perry & Yuval-Greenberg, 2020, 2021). In Exp. 1, we observed that saccades were inhibited to a larger degree when the previous foreperiod matched the current foreperiod (SFD of zero) compared to when the previous foreperiod was longer in duration (negative SFD. The opposite was generally true for trials with shorter previous foreperiod (positive SFD) yet this effect was not symmetrical (see **Figure 2B**) – compared to previous matched trials, saccade probability only slightly decreased or did not significantly differ (see **Table 1**). This raises the question of whether the fixed vs. random effect observed in previous studies stemmed from local sequential information rather than from target probability. In these previous studies, the fixed and random trials were included in different blocks: in the fixed blocks the previous trial always matched the present trial, i.e., had SFD zero; In contrast, in the random block, the previous trial could have been longer, equal, or shorter than the present trial. Since the sequential effect is asymmetrical, averaging trials with positive and negative SFDs (as in the random condition) is expected to lead to a higher saccade rate than trials of zero SFD (as in the fixed condition). It could therefore be hypothesized that the sequential effect is at the basis of the difference in saccade rate between the fixed and random conditions (higher pre-stimulus saccade rate for random relative to fixed) rather than temporal expectation, as was previously suggested.

To examine this hypothesis, we reexamined the fixed vs. random effect reported in two previous studies (Amit et al., 2019; Tal-Perry & Yuval-Greenberg, 2020), while controlling for the previous trial. We compared the probability of performing a saccade prior to target onset between the fixed and the random trials while including only trials in which the previous foreperiod was equal to the current foreperiod. Thus, the n-1 identity was matched between the two conditions. Results were analyzed using a GLMM assuming a binomial family of response, with Condition (fixed / random), Foreperiod (continuous), and the interaction between them as within-subject fixed factors, allowing for a random intercept by subject and random slope for each of the main effects by subject.

Figure 3 depicts the descriptive results of this analysis. As can be observed, we found saccade probability to be significantly lower in the fixed compared to the random condition (*χ*^2^(1) = 6.700, *p* = .01). There was no significant effect for foreperiod (*χ*^2^(1) = 0.916, *p* = .338, *BF*_01_= 77.675), but foreperiod significantly interacted with condition (*χ*^2^(1) = 20.500, *p* < .001), owing to the positive slope in the fixed condition compared to the negative slope in the random condition. This reversal in slopes matches what was observed in Amit et al. (2019) – the decrease in oculomotor inhibition in the fixed condition as foreperiod increases could be explained by the increase in temporal uncertainty, while the increase in oculomotor inhibition in the random condition might be the result of the increasing hazard rate, i.e., the likelihood of an event to occur given that it has yet to occur. Note, however, that the increase in oculomotor inhibition is less steep than the increase in the hazard rate, questioning this interpretation and suggesting the possible involvement of additional factors. Overall, these results are consistent with those found when not controlling for the previous trial foreperiod and therefore suggest that the fixed effect observed in previous studies cannot be explained solely as the result of a sequential effect, but likely reflects target predictability.

**Figure 3.**
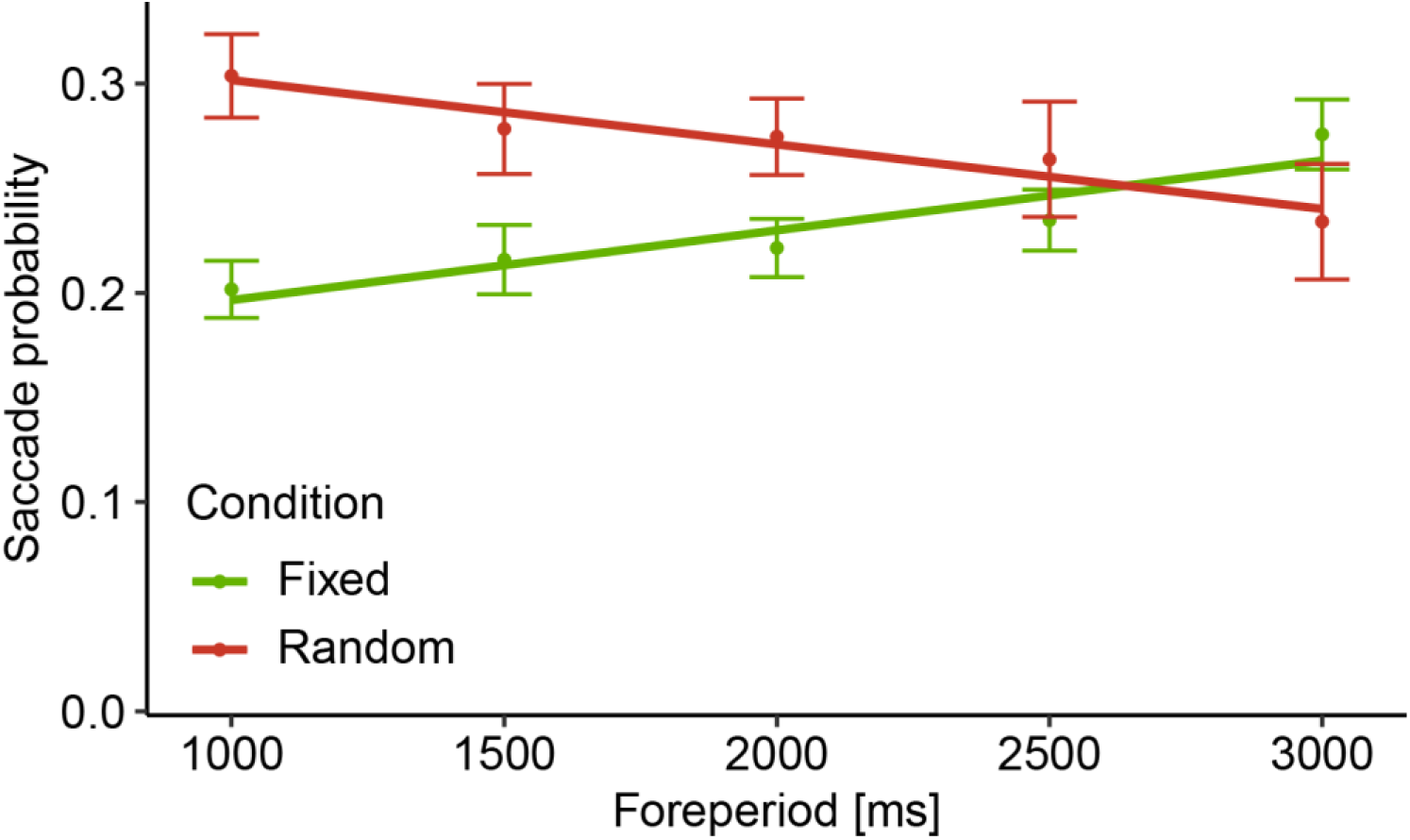
Experiment 2 results. The probability of performing a saccade during the -300 to 0 ms period relative to target onset, as a function of Condition and Foreperiod, with the n-1 trial’s foreperiod matching the current trial’s foreperiod for both conditions. Error bars depict ±1 standard error from the mean, correcting for within-subject variability (Cousineau & O’Brien, 2014). Lines depict linear fit to the observed data. *N* = 40

## Discussion

In this study, we examined whether the pre-stimulus oculomotor inhibition is affected by local sequential information, i.e., the sequential effect – the previous trial history with respect to the current trial. In Exp. 1, we demonstrated that the likelihood of performing a saccade prior to target onset changed as a factor of the relation between the current trial’s foreperiod and the previous trial’s foreperiod, consistent with the typical pattern observed for RT with respect to the sequential effect and demonstrating once more that the pre-stimulus oculomotor inhibition reflects anticipation similarly to RTs.

As a follow-up question, in Exp. 2 we examined whether the sequential effect could provide an alternative explanation to findings observed in previous cue-based temporal expectation studies. In these studies, we found that when the foreperiod was fixed throughout the block, pre-target saccades were inhibited to a larger degree than when the foreperiod varied within the block, for short foreperiods (Abeles et al., 2020; Amit et al., 2019; Badde et al., 2020; Dankner et al., 2017; Tal-Perry & Yuval-Greenberg, 2020, 2021). These findings were interpreted as the result of a higher expectation as target predictability increased. However, in the fixed block of all these cases, the previous trial’s foreperiod always matched the current trial’s foreperiod. Thus, the lower saccade rate found in the fixed condition could be interpreted as resulting from a sequential effect of the previous trial. In Exp. 2, we demonstrated that this interpretation is unlikely – trials from the random condition in which the previous foreperiod matched the current trial’s, nevertheless exhibited a higher saccade rate compared to the fixed condition. These findings are in line with previous RT studies that showed the FP distribution effects cannot be reduced to sequential effects (Los & Agter, 2005; Vallesi et al., 2013). Together, these two experiments indicate that oculomotor inhibition is modulated both by local sequential information and by more global temporal priors, such as target probability, foreperiod distribution and higher-order sequential effects.

### Models of the sequential effect

In Exp. 1, we found an asymmetrical sequential effect of pre-target saccade rate: pre-target saccade rate was higher in trials succeeding trials with longer foreperiod but it was either similar or only slightly lower in trials succeeding a shorter foreperiod. This asymmetry is consistent with the asymmetric sequential effect reported previously with RTs: RTs in trials that follow a trial with longer foreperiod tend to be slower than the RT in trials that follow trials of identical foreperiods, but the opposite is typically not found (e.g., Los et al., 2001; Steinborn & Langner, 2012; Tal-Perry & Yuval-Greenberg, 2022; Vallesi & Shallice, 2007). There are a few theories explaining this asymmetrical pattern in RT data (Los, 2010). We focus here on four of these theories: the *strategic model* (e.g., Alegria & Delhaye-Rembaux, 1975; Niemi & Näätänen, 1981), the *dual-process model* (Vallesi, 2010; Vallesi et al., 2007; Vallesi & Shallice, 2007), and the *trace-conditioning model* (Los et al., 2001), which was later updated to form the *multiple-trace theory* (Los et al., 2014, 2017, 2021; Salet et al., 2022). While these models were developed to explain results in RT data, they may be adapted to interpret the sequential effect observed for the pre-stimulus oculomotor inhibition in the current study.

The *strategic model* (Alegria & Delhaye-Rembaux, 1975; Niemi & Näätänen, 1981) was an initial attempt at explaining the asymmetrical pattern of the sequential effect. According to this view, participants use the target onset time in the *n-1* trial to orient their attention with regard to target onset in the current trial. If time elapses and the target has failed to occur, participants can maintain their preparatory state or shift it toward a later moment when target is likely to occur. Thus, the model predicts RT to be slow when target arrives sooner than anticipated relative to the *n*-1 trial, yet to remain relatively similar if target occurs at the anticipated moment or after it. This theory could explain the asymmetrical sequential effect as demonstrated in Exp. 1 as follows: when the previous foreperiod is shorter than the current foreperiod, participants orient their expectations toward the short period in the current trial, but given that the event has not occurred, they reorient their expectations to the next probable target onset, whose conditional probability is typically higher (e.g., under uniform foreperiod distribution), thus leading to higher expectations and inhibition of eye movements prior to the next target. However, the strategic model was criticized for not providing a full explanation of the sequential effects, particularly their influence in the case of 100% valid cues (Los & Heslenfeld, 2005).

The criticism raised against the strategic model led to the development of the *dual-process model* (Vallesi, 2010; Vallesi et al., 2007; Vallesi & Shallice, 2007). According to the dual-process model, sequential effects stem from two factors: an automatic increase in arousal from the previous trial target (arousal carry-over), along with a controlled or intentional monitoring of conditional probability (hazard rate) that varies during the given trial, akin to the description given by the strategic view. The model posits that the former is the source of the sequential effect, while the latter is the reason the sequential effect is asymmetrical. The model’s identification of the intentional monitoring with the hazard rate function fits the observed difference in asymmetry between the two foreperiod distributions in Exp. 1, as different foreperiod distributions lead to different conditional probabilities. This view is also consistent with our previous studies in which we have demonstrated other effects of conditional probabilities on patterns of pre-stimulus oculomotor inhibition (Abeles et al., 2020; Amit et al., 2019; Tal-Perry & Yuval-Greenberg, 2020). The second process, arousal from the previous trial, affects expectations in the current trial. When preceded by a short trial, arousal tends to be high, and RT is accordingly fast. When preceded by a long trial, the prolonged preparation causes exhaustion of alertness, leading to slower RT in the following trial (Steinborn & Langner, 2012; Vallesi & Shallice, 2007). To provide a similar explanation in the case of pre-stimulus oculomotor inhibition, one has to assume that arousal or readiness to respond is positively correlated with oculomotor rate. In a previous study, we showed that the pre-stimulus oculomotor inhibition is independent of motor readiness (Tal-Perry & Yuval-Greenberg, 2021) and thus response readiness is unlikely to explain the higher saccade probabilities observed for negative SFD trials (longer previous trial) in Exp. 1 of the present study (see Figure 2B). Thus, the dual-process model falls short of explaining the full pattern of results in this study. This is consistent with a study by Capizzi et al., (2015), that tested the prediction of the dual-process models using non-aging distributions with and without catch trials, thereby controlling for the intentional component postulated by the model. In this scenario, the model predicts the sequential effect to be symmetrical, yet results of this study showed an asymmetrical sequential effect. The same study found that these results can be explained by the trace-conditioning model, which we now turn to discuss.

As an alternative, the *trace-conditioning model* (Los et al., 2001) suggests that the asymmetrical sequential effect is the result of a single process – the activation of the weighted memory traces of previous trials starting at cue onset, and decaying as time progresses. This model assumes that each critical moment is associated with a conditioned strength: the higher the conditioned strength associated with a critical moment the higher expectations will be if the target occurs at that moment (Los 2010). However, this model posited that the effect of FP distribution on RT was simply the consequence of the sequential effect, yet the sequential effect was shown to be inadequate in explaining it (Los & Agter, 2005). It was further shown that the sequential effect could be manipulated by changing the inter-trial interval while leaving the FP distribution effect intact (Vallesi et al., 2013), further highlighting that the latter is not a consequence of the former.

The trace-conditioning model formed the basis of a more recent model, the *multiple-trace theory* (Los et al., 2014, 2017, 2021; Salet et al., 2022), according to which the onset of the cue in each trial triggers a motor inhibition which prevents the execution of a premature response during the foreperiod interval. The target onset activates a second neuronal population to elicit a response. This pattern of inhibition followed by activation constitutes the preparatory temporal profile, which is saved as a trace in memory after each trial and can be identified with the expectation process. At each cue onset, the existing memory traces, accumulated over previous trials, are reactivated and are aggregated to a preparatory pattern which determines when will inhibition wane and activation peak within the current trial. Due to the dissipation of memory over time, recent trials contribute more strongly to the activation than older trials. As the preparatory pattern is an aggregation of previous trials, different foreperiod distributions are expected to lead to different preparatory patterns, as the mixture of previous foreperiod (i.e., the higher-order sequential effects) should vary according to distribution. This, in turn, explains the different RT patterns that are induced by different foreperiod distributions.

Given this model, it is clear to see how sequential effects come about. Trials in which the previous foreperiod matches the current foreperiod would lead to a better preparation (more activation and less inhibition) at target onset, compared to trials where there is a mismatch. Trials whose previous foreperiod was longer than the current foreperiod would result in a high level of inhibition around target onset, thereby leading to a slower response. To explain the asymmetry in sequential effects of RT data, the multiple-trace theory postulates that the memory trace builds up and dissipates slowly over the trial. Thus, in trials whose previous foreperiod was shorter than the current foreperiod, the inhibitory content of the previous memory trace would dissipate by target onset, and thus would not contribute to the preparatory profile, meaning that RT would be relatively fast compared to the inverse scenario. Unlike other competing models, The formalized version of this model was shown to make quantitatively-correct predictions of various temporal phenomena, including the sequential effect (Salet et al., 2022).

Can the multiple trace model explain the results observed in our study? Like its predecessor, the trace conditioning model, this model places great importance on the role of inhibition in building up expectations. The findings from this and previous studies on oculomotor inhibition fit with this view – as inhibition builds up to the expected target, saccade rates are lowered, with inhibition being released after target onset. This interpretation fits with the results observed in Exp. 1, with saccade probability being lower when the foreperiod of the previous and current trials matched as compared to where the previous trial was longer in duration (see Figure 2B, negative vs. zero value). The observed asymmetry in pre-stimulus oculomotor inhibition in Exp. 1 (Figure 2B, positive values) can also be explained by the model – in trials where the previous foreperiod was shorter than the current foreperiod, the inhibitory content of the previous memory trace dissipates by target onset, and therefore does not negatively affects expectation, which translates into relatively lower saccade rate. The increase in saccade rate observed for increasing positive SFDs (shorter previous trial) could likewise be accounted for by the dissipation of activation postulated by the model. Lastly, the observed difference in oculomotor inhibition for the uniform and inverse-U-shaped distributions can be explained by differences in the aggregated traces between the two distributions – there are fewer trials with an extreme foreperiod difference in the inverse-U-shaped distribution compared to the uniform distribution, meaning thataggregated activity is predicted to be low at late time points during the trial in the inverse-U-shaped condition, and this translates into a higher saccade probability at extreme positive values as depicted in Figure 2B. Thus, of the presented alternatives, our results are best explained by the multiple trace theory.

## Conclusions

Our results demonstrate that the pre-stimulus oculomotor inhibition is modulated by temporal information that stems from a recent experience. This study joins a growing body of studies that demonstrate that pre-stimulus oculomotor inhibition reflects different types of temporal expectation processes, based on rhythms (Dankner et al., 2017), cue associations (Abeles et al., 2020; Amit et al., 2019; Tal-Perry & Yuval-Greenberg, 2020, 2021) or hazard rate function (Tal-Perry & Yuval-Greenberg, 2020); with the degree of anticipation correlated with the degree of inhibition (Tal-Perry & Yuval-Greenberg, 2020). The present study expands this list by showing that anticipation based on local sequential information is similarly correlated with pre-stimulus oculomotor inhibition. Together, this series of studies demonstrate that whenever there is temporal anticipation, there is also inhibition of eye movements. This study additionally provides a metric to study sequential effects without requiring a response from the participant. This metric may allow studying the sequential effect in uncooperative populations, such as infants and toddlers. An open question that remains following this study is whether the oculomotor sequential effect depends on attention. Future studies could manipulate attention to the target and examine the effect of this manipulation on the oculomotor inhibition effect.

## Data Availability statement

The datasets used in Exp. 1 and 2 and an R-markdown file that reproduces all the reported modeling, statistical analyses, and graphs within the paper are uploaded to the Open Science Foundation repository and are available at doi.org/10.17605/OSF.IO/PV3N2

