## Supplementary material for "Sequential effect and temporal orienting in pre-stimulus oculomotor inhibition"

In Experiment 1, we observed a sequential effect on pre-stimulus oculomotor inhibition. The results of that experiment included saccades of all sizes. Here, we explored whether the same pattern of results holds for saccades of different sizes. Since the vast majority (91.12%) of saccades recorded in this experiment were microsaccades (<1 visual degree), there was an insufficient number of trials to calculate the sequential effect on macrosaccades. Here, we reanalyzed the data from Exp. 1 while focusing solely on microsaccades.

As can be observed in **Figure S1**, the pattern of results closely resembled the pattern observed in Exp. 1 - microsaccades were inhibited to a larger degree when the previous foreperiod matched the current foreperiod, relative to when the previous foreperiod was shorter in duration (negative Standardized Foreperiod Difference, SFD), and this pattern was asymmetrical for positive SFD. As in the main results, this led to a significant effect of SFD ($\chi^2(2) = 32.548$, $p < .001$) with a significant negative linear (log estimate -0.244, $z = -6.240$, $p < .001$) and significant positive quadratic (log estimate 0.080, $z = 4.554$, $p < .001$) components. Here too, the SFD significantly interacted with Foreperiod Distribution ($\chi^2(2) = 14.237$, $p < 0.001$), with distributions differing in both the linear (log estimate 0.034, $z = 2.981$, $p = .003$) and quadratic (log estimate 0.024, $z = 2.596$, $p = .009$) components. Lastly, we again found a significant effect for Foreperiod Distribution ($\chi^2(1) = 48.502$, $p < .001$), such that pre-stimulus microsaccade probability was higher for the inverse-U-shaped distribution.

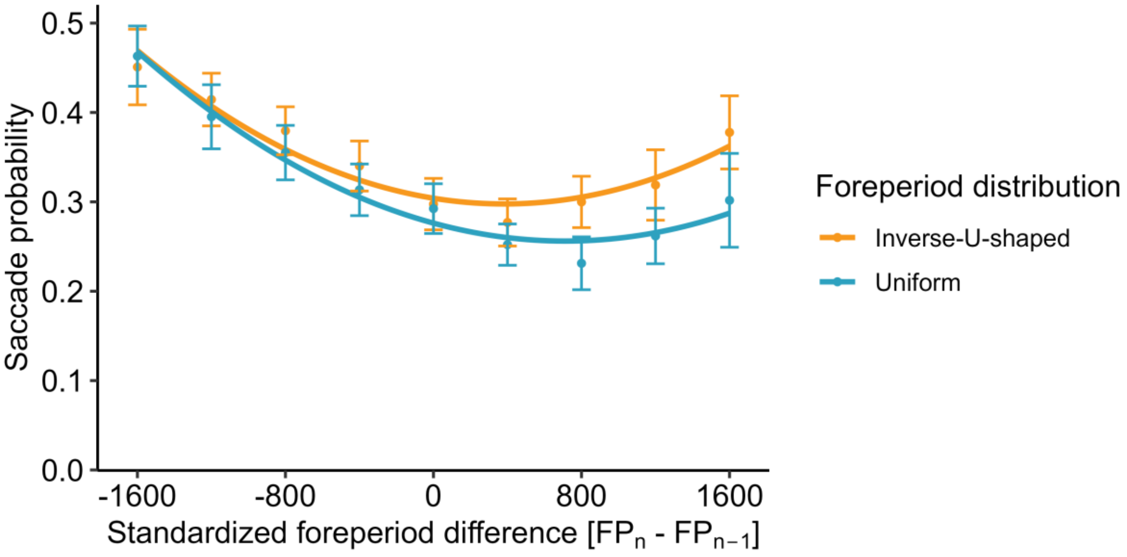


**Figure S1** *Sequential effect on microsaccadic inhibition*. The probability of performing a microsaccade (defined as a saccade of <1 visual degree) during the -300 to 0 ms period relative to target onset, as a function of the difference between the current and previous foreperiod, and the Foreperiod Distribution. Negative values indicate that the previous foreperiod was longer than the current foreperiod, and vice-versa for positive values. Error bars depict ±1 standard error from the mean, correcting for within-subject variability (Cousineau & O'Brien, 2014). Lines depict 2$^{nd}$ polynomial fit to the observed data. $N = 20$ in each distribution.
